## Supplementary figures and images for "3D electron microscopy of the *Leishmania mexicana* cell cycle: Patterns of organelle duplication and segregation and their implications for parasite biology"

### Supplementary Figure 1

Supplementary Figure 1

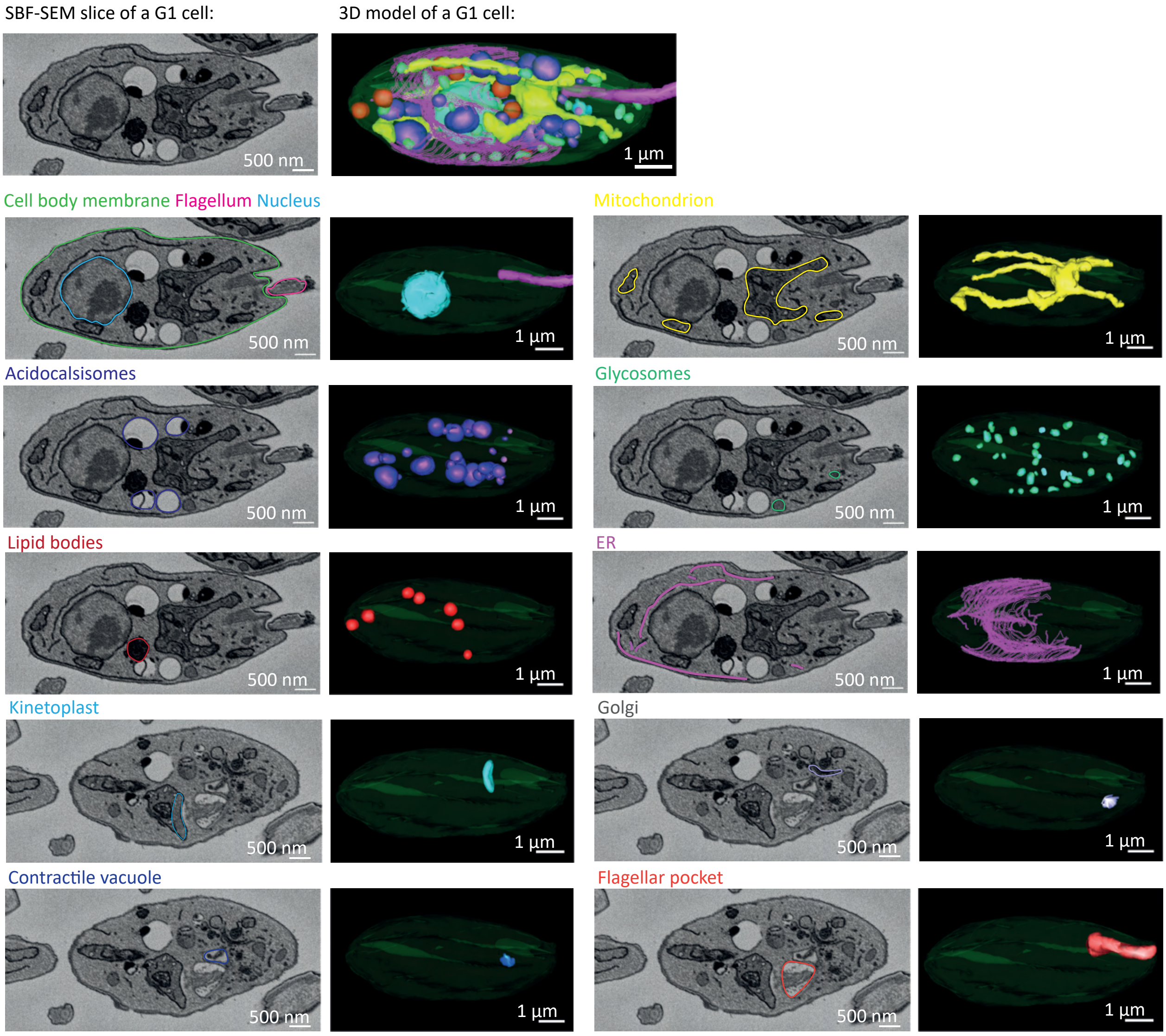

### Supplementary Figure 2

Supplementary Figure 2

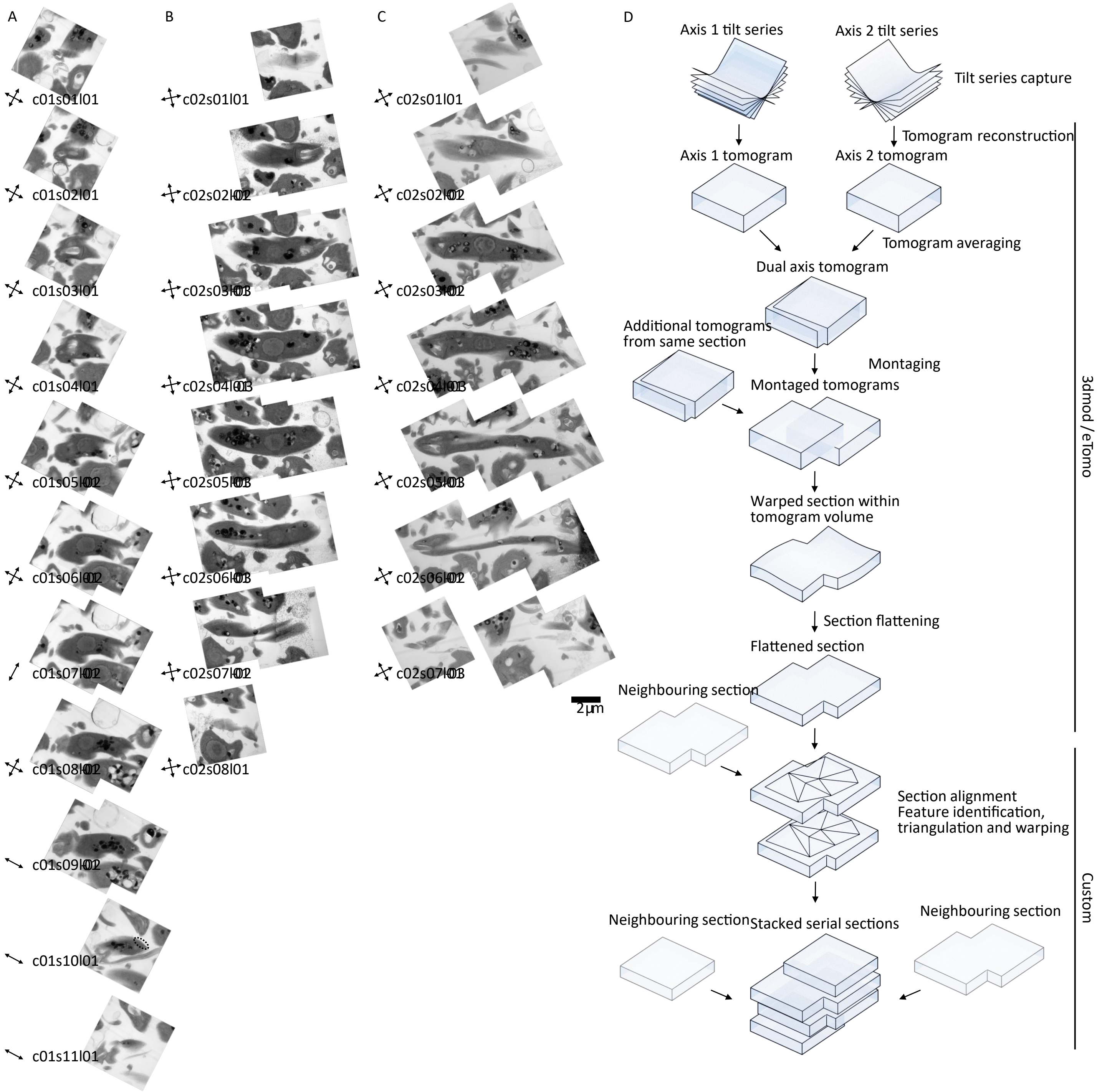
