## Supplementary Table 1 for "3D electron microscopy of the *Leishmania mexicana* cell cycle: Patterns of organelle duplication and segregation and their implications for parasite biology"

**Supplementary Table 1 - culture derived promastigote cells source data**

| | Stage 1 | Stage 2 | Stage 3 | Stage 4 | Stage 5 | Stage 6 | Stage 7 |
|---|---|---|---|---|---|---|---|
| Cell body volume (nm cubed) | 33.61 | 36.53 | 34.39 | 75.65 | 66.97 | 56.9999 | 56.8461 |
| | 36.8 | 59.34 | 54.2 | 70.3 | 82.73 | 62.582 | 77.5187 |
| | 30.87 | 55.13 | 73.96 | 72.1 | 72.93 | 71.2494 | 68.2573 |
| | 45.35 | 48.8 | 38.21 | 53.96 | 72.93 | 69.2543 | 82.1184 |
| | 30.82 | 38.85 | | 59.97 | 58.76 | 69.1612 | |
| | 40.89 | 51.58 | | 39.86 | 77.56 | 81.231 | |
| | | 38.87 | | | | | |
| | | 43.97 | | | | | |
| | | 48.67 | | | | | |
| Cell body surface area (nm squared) | 90.36 | 91.56 | 147.1 | 109.1 | 137.8 | 145.288 | 127.837 |
| | 84.36 | 135.9 | 142.3 | 148.4 | 155.7 | 143.661 | 165.878 |
| | 77.69 | 135.9 | 155.7 | 139.3 | 135.9 | 127.123 | 193.459 |
| | 104.8 | 114.1 | 97.87 | 124.9 | 135.9 | 168.908 | 176.721 |
| | 72.71 | 105.4 | | 128.4 | 136.5 | 78.054 | |
| | 145.5 | 125 | | 112.8 | 157.2 | 171.132 | |
| | | 109.4 | | | | | |
| | | 110.9 | | | | | |
| | | 115.2 | | | | | |

| | Stage 1 | Stage 2 | Stage 3 | Stage 4 | Stage 5 | Stage 6 | Stage 7 - Daughter cell with OF | Stage 7 - Daughter cell with NF |
|---|---|---|---|---|---|---|---|---|
| Cell body length (µm) | 8.37 | 12.3 | 13.48 | 8.112 | 7.32 | 8.03 | 7.55 | 7.41 |
| | 8.11 | 13.18 | 13.03 | 16.9 | 8.83 | 7.99 | 7.08 | 7.69 |
| | 9.63 | 11.38 | 12.69 | 7.7 | 6.96 | 7.48 | 9.13 | 9.15 |
| | 10.28 | 13.71 | 13.5 | 7.322 | 7.22 | 8.92 | 8.07 | 7.7 |
| | 7.15 | 12.5 | | 9.4 | 8.17 | 6.64 | | |
| | 11.12 | 12.48 | | 7.8 | 9.08 | 9.41 | | |
| | | 14.55 | | | | | | |
| | | 12.77 | | | | | | |
| | | 12.12 | | | | | | |

| | Stage 1 | Stage 2 | Stage 3 | Stage 4 | Stage 5 | Stage 6 | Stage 7 |
|---|---|---|---|---|---|---|---|
| Old flagellum (OF) length (µm) | 10.4<br>11<br>11.4<br>18<br>10<br>13.8 | 15<br>16.7<br>14.6<br>15<br>13.3<br>15.3<br>22.3<br>14<br>15.3 | 13.6<br>9<br>17.2<br>11.2 | 10.1<br>10.3<br>21.5<br>12.5<br>11.8<br>15.6 | 14.3<br>22.1<br>18.1<br>13.8<br>19.2<br>22.5 | 29<br>13.4<br>20.9<br>15.24<br>20.18<br>4 | 14.43<br>15.85<br>14.09<br>17.3 |
| New flagellum (NF) length (µm) | | 0.597<br>0.335<br>1.9<br>0.702<br>0.281<br>0.205<br>1.529<br>1.3<br>0.832 | 5.1<br>6.2<br>3.63<br>3.9 | 4.8<br>3.7<br>5.5<br>9.9<br>8.2<br>8.4 | 4<br>8.3<br>6.5<br>6.8<br>8.2<br>7.38 | 8.34<br>7<br>9.98<br>10.3<br>6.58<br>10.8 | 10.2<br>10.59<br>10.06<br>8.5 |

| | Stage 1 | Stage 2 | Stage 3 | Stage 4 | Stage 5 | Stage 6 - Daughter cell with OF | Stage 6 - Daughter cell with NF | Stage 7 - Daughter cell with OF | Stage 7 - Daughter cell with NF |
|---|---|---|---|---|---|---|---|---|---|
| Nucleus volume (nm cubed) | 2.32<br>2.19<br>3.04<br>3.13<br>2.44<br>3.36 | 2.68<br>3.73<br>4.38<br>4.28<br>3.03<br>4.82<br>3.66<br>4.55<br>4.34 | 3.54<br>7.09<br>6.13<br>4.54 | 5.07<br>3.63<br>5.92<br>3.86<br>4.78<br>3.09 | 5.92<br>6.14<br>5.99<br>6.71<br>4.86<br>6.09 | 2.24<br>2.03<br>2.61<br>1.17<br>2.42<br>2.19 | 2.36<br>1.92<br>2.39<br>0.903<br>2.27<br>2.43 | 2.43<br>2.23<br>2.12<br>2.84 | 2.37<br>2.31<br>2.5<br>2.88 |
| Nucleus surface area (nm squared) | 10.06<br>9.58<br>11.15<br>11.59<br>9.93<br>12.29 | 10.39<br>12.85<br>14.40<br>14.06<br>11.45<br>15.60<br>12.97<br>14.85<br>14.17 | 13.15<br>20.95<br>19.50<br>13.31 | 16.76<br>15.64<br>23.42<br>18.27<br>18.01<br>15.76 | 22.95<br>26.26<br>25.04<br>23.71<br>21.33<br>21.53 | 9.28<br>9.38<br>10.75<br>6.34<br>10.29<br>9.58 | 9.75<br>9.36<br>10.11<br>4.87<br>9.75<br>10.65 | 9.88<br>9.77<br>9.14<br>12.16 | 9.72<br>10.02<br>10.38<br>11.94 |

| | Stage 1 | Stage 2 | Stage 3 | Stage 4 | Stage 5 | Stage 6 - Daughter cell with OF | Stage 6 - Daughter cell with NF | Stage 7 - Daughter cell with OF | Stage 7 - Daughter cell with NF |
|---|---|---|---|---|---|---|---|---|---|
| Number of nuclear pores | 27<br>30<br>27<br>25<br>32<br>22 | 36<br>42<br>45<br>40<br>36<br>41<br>42<br>41<br>40 | 44<br>67<br>31<br>39 | 59<br>38<br>42<br>49<br>47<br>39 | 35<br>41<br>28<br>66<br>48<br>31 | 19<br>39<br>26<br>24<br>40<br>22 | 27<br>30<br>29<br>22<br>24<br>19 | 24<br>34<br>38<br>42 | 25<br>39<br>33<br>48 |

| | Stage 1 | Stage 2 | Stage 3 | Stage 4 | Stage 5 - Daughter cell with OF | Stage 5 - Daughter cell with NF | Stage 6 - Daughter cell with OF | Stage 6 - Daughter cell with NF | Stage 7 - Daughter cell with OF | Stage 7 - Daughter cell with NF |
|---|---|---|---|---|---|---|---|---|---|---|
| Kinetoplast membrane volume (µm cubed) | 0.15<br>0.11<br>0.14<br>0.23<br>0.15<br>0.18 | 0.23<br>0.29<br>0.28<br>0.24<br>0.21<br>0.16<br>0.22<br>0.40<br>0.26 | 0.28<br>0.40<br>0.28<br>0.25 | 0.22<br>0.43<br>0.43<br>0.36<br>0.21<br>0.22 | 0.15<br>0.20<br>0.22<br>0.20<br>0.18<br>0.14 | 0.19<br>0.16<br>0.24<br>0.13<br>0.13<br>0.12 | 0.19<br>0.15<br>0.15<br>0.12<br>0.11 | 0.12<br>0.20<br>0.11<br>0.11<br>0.10 | 0.15<br>0.15<br>0.11<br>0.14 | 0.12<br>0.11<br>0.11<br>0.13 |

| | Stage 1 | Stage 2 | Stage 3 | Stage 4 | Stage 5 | Stage 6 | Stage 6 - Daughter cell with OF | Stage 6 - Daughter cell with NF | Stage 7 - Daughter cell with OF | Stage 7 - Daughter cell with NF |
|---|---|---|---|---|---|---|---|---|---|---|
| Mitochondrion volume (µm cubed) | 2.33<br>2.38<br>1.92<br>3.10<br>1.78<br>2.62 | 3.01<br>4.14<br>3.34<br>3.23<br>2.48<br>2.63<br>2.74<br>3.02<br>3.45 | 3.15<br>4.61<br>4.75<br>2.92 | 4.48<br>3.45<br>5.01<br>4.13<br>3.58<br>3.41 | 3.67<br>4.99<br>4.18<br>4.89<br>3.99<br>5.46 | 3.98<br>4.72<br>5.37 | 1.47<br>2.85<br>1.94 | 1.69<br>1.92<br>1.98 | 2.83<br>2.39<br>2.54<br>2.58 | 2.10<br>2.39<br>2.40<br>3.00 |
| Mitochondrion surface area (µm squared) | 27.42<br>25.70<br>34.50<br>49.20<br>32.65<br>45.21 | 43.95<br>60.95<br>52.07<br>46.84<br>39.09<br>51.76<br>47.66<br>38.85<br>47.51 | 44.06<br>77.56<br>76.04<br>40.61 | 49.75<br>57.63<br>74.00<br>57.32<br>44.92<br>32.14 | 61.29<br>71.71<br>65.16<br>83.21<br>69.84<br>74.34 | 47.30<br>70.90<br>57.80 | 26.60<br>30.82<br>12.20 | 30.14<br>11.13<br>17.10 | 36.09<br>26.44<br>28.25<br>31.99 | 33.11<br>25.93<br>27.49<br>35.95 |

| | Stage 1 | Stage 2 | Stage 3 | Stage 4 | Stage 5 | Stage 6 - Daughter cell with OF | Stage 6 - Daughter cell with NF | Stage 7 - Daughter cell with OF | Stage 7 - Daughter cell with NF |
|---|---|---|---|---|---|---|---|---|---|
| Number of lipid bodies per cell | 3<br>13<br>5<br>5<br>4<br>4 | 4<br>5<br>5<br>6<br>1<br>12<br>5<br>7<br>5 | 10<br>6<br>9<br>13 | 12<br>18<br>7<br>7<br>9<br>11 | 12<br>5<br>10<br>6<br>12<br>3 | 7<br>4<br>2<br>5<br>6<br>7 | 3<br>1<br>5<br>6<br>5<br>5 | 2<br>6<br>5<br>4 | 3<br>8<br>5<br>13 |
| Total volume of lipid bodies per cell | 0.16<br>0.41<br>0.36<br>0.35<br>0.03<br>0.48 | 0.23<br>0.38<br>0.16<br>0.21<br>0.00<br>0.16<br>0.31<br>0.26<br>0.22 | 0.50<br>0.50<br>0.43<br>0.59 | 0.40<br>0.87<br>0.66<br>0.18<br>0.43<br>0.58 | 0.49<br>0.46<br>0.41<br>0.24<br>0.80<br>0.22 | 0.52<br>0.42<br>0.16<br>0.14<br>0.02<br>0.62 | 0.22<br>0.13<br>0.25<br>0.30<br>0.01<br>0.21 | 0.07<br>0.14<br>0.27<br>0.23 | 0.15<br>0.30<br>0.30<br>0.64 |

| | Stage 1 | Stage 2 | Stage 3 | Stage 4 | Stage 5 | Stage 6 - Daughter cell with OF | Stage 6 - Daughter cell with NF | Stage 7 - Daughter cell with OF | Stage 7 - Daughter cell with NF |
|---|---|---|---|---|---|---|---|---|---|
| Number of acidocalcisomes per cell | 36<br>20<br>53<br>63<br>65<br>41 | 41<br>59<br>61<br>44<br>52<br>64<br>71<br>65<br>41 | 58<br>81<br>48<br>61 | 119<br>79<br>72<br>110<br>75<br>78 | 75<br>97<br>47<br>60<br>80<br>69 | 21<br>31<br>35<br>53<br>53<br>54 | 24<br>24<br>32<br>38<br>37<br>42 | 37<br>29<br>54<br>54 | 32<br>28<br>35<br>40 |
| Total volume of acidocalcisomes per cell | 2.26<br>3.06<br>1.62<br>2.54<br>1.55<br>4.57 | 2.14<br>3.19<br>2.46<br>3.21<br>2.34<br>1.64<br>1.05<br>1.80<br>3.26 | 1.48<br>4.95<br>5.66<br>1.19 | 3.96<br>2.55<br>5.22<br>2.02<br>5.94<br>1.71 | 3.51<br>6.79<br>6.09<br>3.86<br>2.15<br>3.89 | 3.86<br>4.77<br>1.21<br>1.52<br>1.50<br>2.99 | 4.41<br>3.69<br>1.77<br>2.74<br>1.52<br>2.74 | 1.79<br>4.20<br>3.17<br>3.88 | 2.06<br>5.26<br>3.37<br>4.56 |

| | Stage 1 | Stage 2 | Stage 3 | Stage 4 | Stage 5 | Stage 6 - Daughter cell with OF | Stage 6 - Daughter cell with NF | Stage 7 - Daughter cell with OF | Stage 7 - Daughter cell with NF |
|---|---|---|---|---|---|---|---|---|---|
| Number of glycosomes per cell | 40<br>40<br>24<br>45<br>28<br>45 | 33<br>71<br>42<br>45<br>44<br>57<br>50<br>31<br>41 | 20<br>78<br>71<br>29 | 60<br>76<br>71<br>58<br>62<br>50 | 55<br>81<br>65<br>62<br>72<br>67 | 34<br>42<br>31<br>48<br>29<br>56 | 26<br>29<br>33<br>30<br>26<br>38 | 24<br>28<br>38<br>36 | 21<br>35<br>31<br>37 |
| Total volume of glycosomes per cell | 0.32<br>0.34<br>0.20<br>0.37<br>0.22<br>0.29 | 0.32<br>0.49<br>0.28<br>0.28<br>0.26<br>0.22<br>0.17<br>0.27<br>0.25 | 0.30<br>0.66<br>0.51<br>0.36 | 0.59<br>0.36<br>0.56<br>0.56<br>0.42<br>0.36 | 0.38<br>0.49<br>0.38<br>0.53<br>0.46<br>0.36 | 0.30<br>0.31<br>0.15<br>0.42<br>0.01<br>0.32 | 0.23<br>0.25<br>0.21<br>0.35<br>0.02<br>0.27 | 0.25<br>0.33<br>0.30<br>0.26 | 0.22<br>0.34<br>0.27<br>0.31 |

**Sand fly derived promastigote cells source data**

| | Cell 1 | Cell 2 | Cell 3 | Cell 4 | Cell 5 | Cell 6 | Cell 7 | Cell 8 | Cell 9 | Cell 10 |
|---|---|---|---|---|---|---|---|---|---|---|
| Flagellum length (µm) | 12.9 | 12.32 | 15 | 12.5 | 8.5 | 11.26 | 9.8 | 21 | 5.4 | 5 |
| Cell body length (µm) | 8 | 6 | 12.3 | 11.9 | 9.6 | 7.7 | 13.2 | 12.6 | 5.6 | 6.8 |
| Cell body volume (µm) | 9.03 | 9.09 | 9.43 | 8.78 | 8.76 | 5.92 | 7.99 | 13.48 | 11.64 | 19.54 |
| Cell body SA (µm squared) | 39.76 | 34.39 | 39.80 | 43.97 | 43.13 | 27.30 | 40.91 | 58.26 | 36.21 | 59.2 |
| Nucleus volume (µm cubed) | 1.09 | 1.11 | 0.94 | 0.99 | 1.01 | 0.89 | 0.93 | 1.23 | 1.11 | 1.61 |

| | Cell 1 | Cell 2 | Cell 3 | Cell 4 | Cell 5 | Cell 6 | Cell 7 | Cell 8 | Cell 9 | Cell 10 |
|---|---|---|---|---|---|---|---|---|---|---|
| Number of acidocalcisomes | 54 | 37 | 30 | 26 | 23 | 26 | 20 | 41 | 26 | 69 |
| Acidocalcisomes total volume (µm cubed) | 0.06 | 0.13 | 0.08 | 0.20 | 0.10 | 0.05 | 0.02 | 0.07 | 0.06 | 0.23 |
| Number of lipid bodies | 6 | 12 | 15 | 18 | 19 | 9 | 24 | 15 | 14 | 21 |
| Lipid bodies total volume (µm cubed) | 0.04 | 0.67 | 0.73 | 0.38 | 0.40 | 0.27 | 0.21 | 1.08 | 0.63 | 1.55 |
